# Developing High Content Imaging Functional Panels to Characterize Synovial Fibroblasts in Rheumatoid Arthritis

**DOI:** 10.64898/2026.06.23.733987

**Authors:** Phatthamon Laphanuwat, Ege Ezen, Christof Seiler, Caroline Ospelt

## Abstract

**Objective:** To develop and apply a preclinical functional imaging assay for visualizing and analyzing activated synovial fibroblasts (SFs) at single-cell resolution using high-content imaging.

**Methods:** A multiparametric functional imaging assay was developed to simultaneously interrogate six key cellular processes in cultured SFs from non-inflammatory control (NIC), osteoarthritis (OA) and rheumatoid arthritis (RA) patients. Two complementary fluorescent panels — comprising LipidTOX, MitoSOX, TMRM, EdU Click-iT, CYTO-ID, and Sir-Lysosome — collectively captured autophagy dynamics, mitochondrial health, lipid metabolism, and cellular proliferation within a single imaging workflow. Automated image acquisition and quantitative feature extraction via CellProfiler yielded approximately 1,200 morphological and intensity-based features per cell, enabling high-dimensional, unbiased phenotypic profiling at the individual cell level.

**Results:** Application of this assay revealed marked heterogeneity in basal cellular functions among SFs stratified by disease state, and robustly differentiated between NIC, OA and RA SFs. Stimulation with inflammatory cytokines (TNF-α, IL-1β, IFNƔ) and toll-like receptor ligands (LPS, poly I:C) elicited distinct, stimulus-dependent phenotypic responses across disease groups. Multiparametric analysis and feature importance ranking identified IL-1β as a key driver of enhanced autophagic activity, accompanied by significant remodeling of lipid metabolic profiles.

**Conclusion:** We developed a scalable, sensitive approach for dissecting functional heterogeneity in primary SF cultures, revealing previously unappreciated complexity in SF biology across disease states. Our approach provides a robust framework for high-throughput drug screening and identification of candidate therapeutics selectively targeting pathogenic fibroblast functions in inflammatory arthritis.

## Introduction

Several lines of evidence support synovial fibroblasts (SFs) as therapeutic targets in rheumatoid arthritis (RA). They expand drastically and are at the forefront of the synovial tissue that invades and erodes cartilage and bone (1–4). SFs were shown to be activated by pro-inflammatory cytokines — particularly TNF-α, IL-1β, and IFNƔ— and by endogenous TLR ligands released during tissue damage, i.e. damage-associated molecular patterns (DAMPs) (5,6). Activated SFs resist apoptosis and secrete matrix-degrading molecules (e.g. MMP1 and MMP3) as well as pro-inflammatory cytokines, in particular IL-6. Intriguingly, they maintain an invasive, pro-inflammatory phenotype *in vitro* which can be leveraged for *in vitro* drug discovery (7). Furthermore, treatment resistance in RA has been linked to a distinct stromal-fibroblast gene signature, with enrichment of extracellular matrix components and fibroblast-related signaling modules defining a fibroid, pauci-immune synovial pathotype were associated with poor response to both conventional and biologic therapies (8,9).

The activation of SFs in RA involves profound metabolic reprogramming including changes in autophagic flux, mitochondrial dysfunction, oxidative stress, and dysregulated lipid metabolism, which collectively sustain the hyperproliferative, and invasive SF phenotype (10–12). These metabolic hallmarks represent both mechanistic drivers of RA pathology and underexploited therapeutic vulnerabilities. Current therapeutic strategies do not directly target SF modulation and therapies that normalize these functions of SF in RA would be highly relevant.

High-content screening (HCS) combines automated multi-channel fluorescence imaging with quantitative feature-based analysis across hundreds of cellular parameters simultaneously (13). Translating these raw fluorescence images into biologically meaningful measurements demands an algorithm that is both comprehensive and reproducible across multiple subcellular compartments. CellProfiler is an open-source tool for quantitative image analysis, that has been widely applied in drug discovery, and phenotypic cell screening (14,15). Its modular structure enables systematic extraction of ∼1,200 features per cell including measures of morphology, texture, intensity, and spatial organization, and integrates directly with downstream machine learning for high-dimensional phenotypic profiling. However, preclinical drug discovery assays, which are primarily used in cancer research, typically rely on single-readout, population-averaged assays that fail to capture the complex functional changes of RA SFs, which is a significant translational gap to identify fibroblast-targeting therapies.

Here, we developed and validated a novel multiparametric HCS assay applied to cultured primary SFs, using two complementary fluorescent panels to profile autophagy, mitochondrial health, lipid metabolism, and proliferation. Using this approach, we were able to distinguish cultured SFs derived from non-inflammatory controls (NIC), osteoarthritis (OA), and RA patients, and to characterize stimulus-dependent functional responses. In particular, we identified functional and morphological differences between ‘inflammatory’, IL-1β-induced autophagy and ‘classical’ rapamycin-induced autophagy. Thus, our approach provides a scalable, biologically comprehensive framework for capturing and characterizing SF activation *in vitro* and can in future be used to identify candidate therapeutics in inflammatory arthritis.

## Methods

### Primary cell lines

Synovial tissue was obtained from patients undergoing joint replacement surgery or ultrasound-guided fine-needle biopsies, with written informed consent and local Swiss ethics committee approval (KEK 2019_00674, 2021_00092). RA patients fulfilled the 2010 ACR/EULAR classification criteria. Non-inflammatory controls (NIC) were derived from patients without inflammatory joint disease undergoing elective arthroscopic procedures and who signed a informed consent form (Ethic approval. KEK 2021_01818). All SF were used under the ethic license KEK 2019_00115. SFs were isolated and cultured in DMEM (Gibco®) supplemented with 10% FCS, 2 mM L-glutamine, 50 U/mL penicillin, 50 µg/mL streptomycin, 10 mM HEPES, and 0.2% amphotericin B at 37°C with 5% CO₂. For experiments, fibroblasts were seeded at 6,000 cells/well in 96-well plates.

### Chemicals and drugs

For panel validation, cells were treated with positive control compounds for 24 hours at 37°C: MG-132 (10 µM; Sigma, M7449) for proteasome inhibition, and rapamycin (0.5 µM; Enzo, ENZ-51031) combined with chloroquine (10 µM; Enzo, ENZ-51031) for autophagy induction. Verapamil (10 µM; Spirochrome, SC012) was included in Panel 1 as an efflux pump inhibitor to enhance SiR-Lysosome staining.

For inflammatory stimulation, fibroblasts were treated for 24 hours prior to staining with: TNFα (10 ng/mL; R&D Systems), IL-1β (1 ng/mL; R&D Systems), IFNƔ-γ (10 ng/mL; ImmunoTools), Poly I:C (5 µg/mL; Invitrogen), and LPS (100 ng/mL; Abbexa).

### Functional staining

Refer to Supplementary Methods.

### Imaging

Images were acquired on a CellInsight CX7 High Content Screening platform (Thermo Fisher Scientific) using HCS Studio software, with single-channel images exported as TIFF files. Panel 1 was imaged at 20× across 36 fields per well in four fluorescence channels: Panel 2 at 10× across 4 fields per well in four channels.

### Image analysis, data transformation and normalization

Image analysis was performed using CellProfiler v4.2.1 (42). Images were corrected for uneven illumination prior to segmentation. Nuclei were segmented from Hoechst-stained images using Minimum Cross-Entropy global thresholding, and whole-cell masks were generated by watershed-gradient propagation from nuclear objects using LipidTOX (Panel 1) or TMRM (Panel 2) channels, selected for their broad cytoplasmic distribution and signal intensity. Cells at image borders or with segmentation artifacts or abnormal intensity profiles were excluded.

Data transformation and normalization were performed using a custom R pipeline implemented in the cellpaintr package (http://github.com/christofseiler/cellpaintr), applied independently per plate. Briefly, cells with missing feature values were removed, and images were filtered by cell count (Panel 1: 5–100 cells/image; Panel 2: 30–300 cells/image). Single-cell outliers were excluded based on total feature intensity using perCellQCMetrics (≥3 median absolute deviations from median). Features with zero variance or zero-inflated distributions were removed and remaining non-negative features were log-transformed using robust scaling. After plate-wise normalization, only features common across all plates were retained and merged into a single matrix for downstream analysis. Single-cell features were then aggregated to the image level by averaging across all cells per image. Cell counts per image were visually assessed across batches to confirm consistency.

### Classification using machine learning

Leave-one-patient-out (LOO) cross-validation was used to evaluate classification performance of cell-level features across disease groups within control-treated samples. A random forest classifier was trained iteratively, with cells from one patient held out as the test set and the remaining patients used for training, repeated for each patient to obtain unbiased predictions. Classification probabilities were predicted for each cell using transformed feature matrices as input. ROC curves and LOO diagnostic plots were generated to assess discriminative power, with patient, disease, and plate included as grouping variables. This approach accounts for patient heterogeneity and minimizes overfitting to individual samples.

### Volcano Plots

LOO classification scores were summarized using volcano plots, where each point represents a feature set. For each feature set, scores from treated and untreated control patients were compared using a two-sample t-test (null hypothesis: equal means), with Bonferroni-corrected significance threshold of p < 0.01. The x-axis displays −log10(p-value) and the y-axis displays log2 fold change, calculated as the ratio of mean treatment to mean control scores. A log2 fold change of 1 indicates that the mean treatment score is twice that of the control. Notably, these volcano plots are asymmetric with strictly positive log2 fold changes, as the classification score represents the predicted probability of belonging to the treatment group — a negative fold change would imply below-chance classifier performance.

### Feature importance analysis

This analysis follows the methodology described in Haghighi M *et al*. (43). CellProfiler extracted approximately 1,200 features per panel, with multicollinearity addressed through a two-step feature selection procedure. First, features were clustered across a range of Pearson correlation thresholds and cluster medoids selected as representative variables. Variance inflation factors (VIF) were then computed on each medoid set, and the configuration retaining the most features with VIF < 10 was selected as the optimal reduced panel. Data were subset to the target comparison groups (e.g., control vs. IL-1Β or between disease groups), with optional marker- or measurement-type-specific filtering applied where warranted — for example, restricting the importance analysis to intensity-based features or to features derived from a specific channel such as TMRM. A random forest classifier (1,000 trees) was trained on the reduced feature set, with variable importance quantified as Mean Decrease in Accuracy.

### Autophagic gene analysis

The autophagy gene list was obtained from HADb (24). Bulk RNA-seq data from cytokine-stimulated SFs (23) were DESeq2-normalised, excluding transcripts with fewer than 10 reads. Genes ranked by signed −log10(p) were used as GSEA input (clusterProfiler; 10,000 permutations) against the autophagy gene set. Leading-edge analysis of the IL-1β GSEA identified 25 driver genes, of which 12 directly implicated in autophagosome formation and dynamics were plotted by log2 fold change and −log10(padj).

## Results

### Establishment of Multiplexed Functional Readouts of Metabolism and Stress in SFs

We evaluated six fluorescent dyes targeting key cellular processes in cultured NIC SFs: neutral lipid metabolism (LipidTOX), mitochondrial health and stress (TMRM and MitoSOX), proliferation (EdU Click), and autophagy dynamics, including autophagosomes (CYTO-ID) and lysosomes (Sir-Lysosome, hereafter referred to as “Lysosome”). Considering dye compatibility and limitations, we organized these markers into two distinct panels: Panel 1 comprised LipidTOX, MitoSOX, and EdU (Supplementary Fig. 1A); Panel 2 included CYTO-ID, TMRM, and Lysosome (Supplementary Fig. 1B); see Table 1 (<u>Supplementary Methods</u>) for detailed dye properties including wavelength, filters, fixation compatibility, and live staining capability).

We validated the assays in panel 1 using MG132, a proteasome inhibitor known to induce global cellular stress and halt proliferation. After 24 hours of MG132 treatment, we detected significantly decreased cellular proliferation of RA SFs (Supplementary Fig. 1C). Concurrently, LipidTOX and MitoSOX intensities increased (Supplementary Fig. 1D–E), indicating lipid accumulation and mitochondrial reactive oxygen species generation despite proliferative arrest. Notably, LipidTOX and MitoSOX intensities strongly correlated in unstimulated conditions and under MG132 treatment, highlighting a strong link between lipid metabolism and mitochondrial stress (Supplementary Fig. 1F).

In Panel 2, MG132 treatment increased TMRM signal, reflecting elevated mitochondrial membrane potential (Supplementary Fig. 1G). Autophagy was induced using rapamycin and chloroquine (R+C), which resulted in a marked increase in CYTO-ID signal (Supplementary Fig. 1H) and a decrease in lysosome signal (Supplementary Fig. 1I) after 24 hours, illustrating autophagosome formation and lysosome consumption during autophagy. CYTO-ID and lysosome signals displayed strong correlation following R+C treatment but not in resting state, where autophagosomes were scarcely detected (Supplementary Fig. 1J).

These results demonstrate the utility of our dual-panel assay platform in capturing multiple functional phenotypes of SFs at single-cell resolution, supporting its potential for high-content functional analyses of SFs.

### Fluorescence Intensity Measures Distinguish RA from OA SFs

Differences between cultured SFs from RA and OA patients have been demonstrated at the epigenetic, transcriptional and functional levels, and RA SFs retain a disease-specific invasive phenotype in cell culture over multiple passages (16). We thus wanted to explore whether we can use high content imaging to differentiate between cultured SFs isolated from NIC, OA, and RA donors. We first focused our analysis on fluorescence intensity measures. Mean fluorescence intensity (MFI), which is the most widely used metric for quantifying marker expression, normalizes total signal by cell area. Integrated intensity instead measures total pixel sum within the cell area. Two cells with equivalent MFI but different sizes can carry different total organelle content or marker load. These differences are captured by integrated intensity but obscured by MFI. As first assay, we used CellPainting (17), an established multiplex fluorescence staining approach that captures broad cellular and subcellular morphology. None of the captured morphological features differed between NIC, OA and RA SFs (Fig. 1A, Supplementary Fig. 2A).

**Fig. 1.**
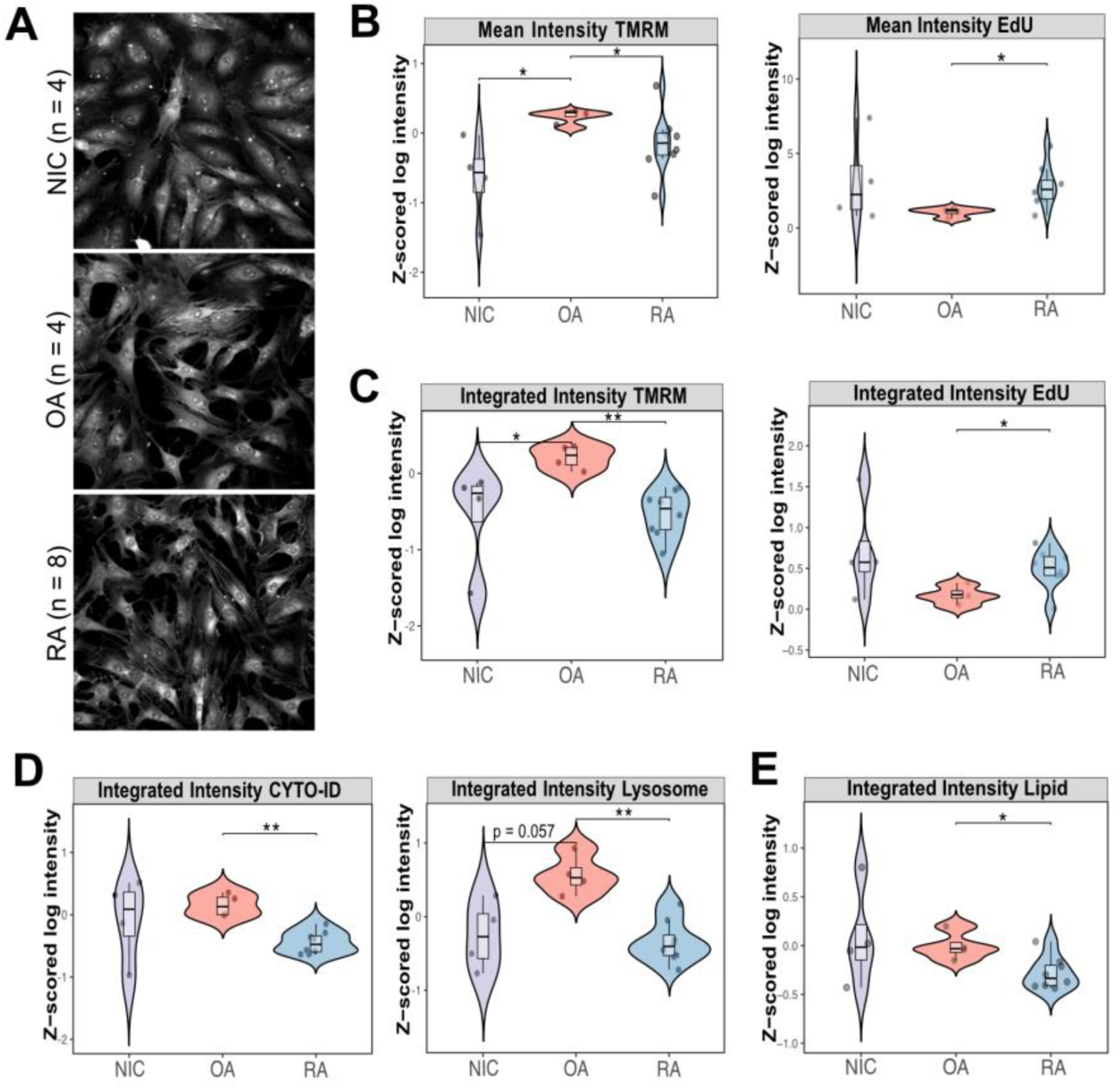
Disease differences based on functional panel simple intensity metrics. Initial evaluation used mean and integrated fluorescence intensities to confirm staining quality and biological relevance at basal conditions. (A) Representative Cell Painting images show comparable fibroblast morphologies across groups; total sample numbers are indicated in parentheses. (B–E) Violin plots comparing NIC (n = 4), OA (n = 4), and RA (n = 8) fibroblasts at baseline: (B) mean fluorescence intensity of TMRM and EdU; (C) integrated intensity of TMRM and EdU; (D) CYTO-ID and Lysosome; (E) neutral lipid. Each dot represents an individual patient. Statistical comparisons used pairwise Wilcoxon tests and Bonferroni-adjusted p-values (** p.adj ≤ 0.01, * p.adj ≤ 0.05).

We then determined whether our functional panel could capture specific pathogenic characteristics of RA SFs. In comparisons across NIC, OA, and RA SFs, MFI-based findings (Fig. 1B) as well as integrated intensity (Fig. 1C) showed higher mitochondrial membrane potential (TMRM) in OA than in RA SFs, but higher proliferative activity (EdU) in RA compared to OA SFs. NIC samples exhibited considerable inter-individual variability, which collectively limited statistical power in this group. There were no significant differences in other MFI comparisons (Supplementary Fig. 2B). Integrated intensity analysis additionally showed higher CYTO-ID and lysosomal content in OA relative to RA SFs (Fig. 1D). This appears to contrast with published reports of elevated autophagy in RA (12,18–20). However, our measurement was done in the resting, unstimulated state, where autophagic activity was generally low (see Supplementary Fig. 1H–J) but could indicate a disturbed basal autophagic flux in RA SFs. Consistent with previous findings (10), RA SFs exhibited lower neutral lipid accumulation than OA SFs as measured by LipidTOX integrated intensity (Fig. 1E).

### Multiparametric Feature Analysis Reveals Distinct Functional Signatures Across Disease States

After evaluating individual features, we applied random forest classifiers with leave-one-out cross-validation to assess whether multidimensional feature relationships could improve disease group discrimination. We defined function-specific feature sets for each marker (e.g., Lipid, EdU etc.), incorporating all relevant feature types associated with that marker, including intensity, texture, and spatial features (see Feature Importance Analysis in Methods).

In multidimensional feature analysis, OA fibroblasts were distinguished from NIC primarily by proliferation (EdU), and mitochondrial stress (MitoSOX) (Fig. 2A), while RA fibroblasts diverged from NIC through lysosomal content and autophagosome (CYTO-ID) (Fig. 2B). Direct OA vs. RA comparison identified lipid metabolism alongside lysosomal, autophagic features and mitochondria membrane potential as most discriminative between the two states (Fig. 2C). Integrating all features within each panel further enhanced discrimination. Panel 1 achieved perfect NIC vs. OA separation but performed poorly for NIC vs. RA, indicating that the metabolism-focused readouts in panel 1 better captured OA-than RA-specific biology (Fig. 2D). Conversely, Panel 2 showed marginal NIC vs. OA discrimination but considerably better NIC vs. RA performance, confirming the added value of autophagy and mitochondrial membrane potential readouts for RA (Fig. 2E).

**Fig. 2.**
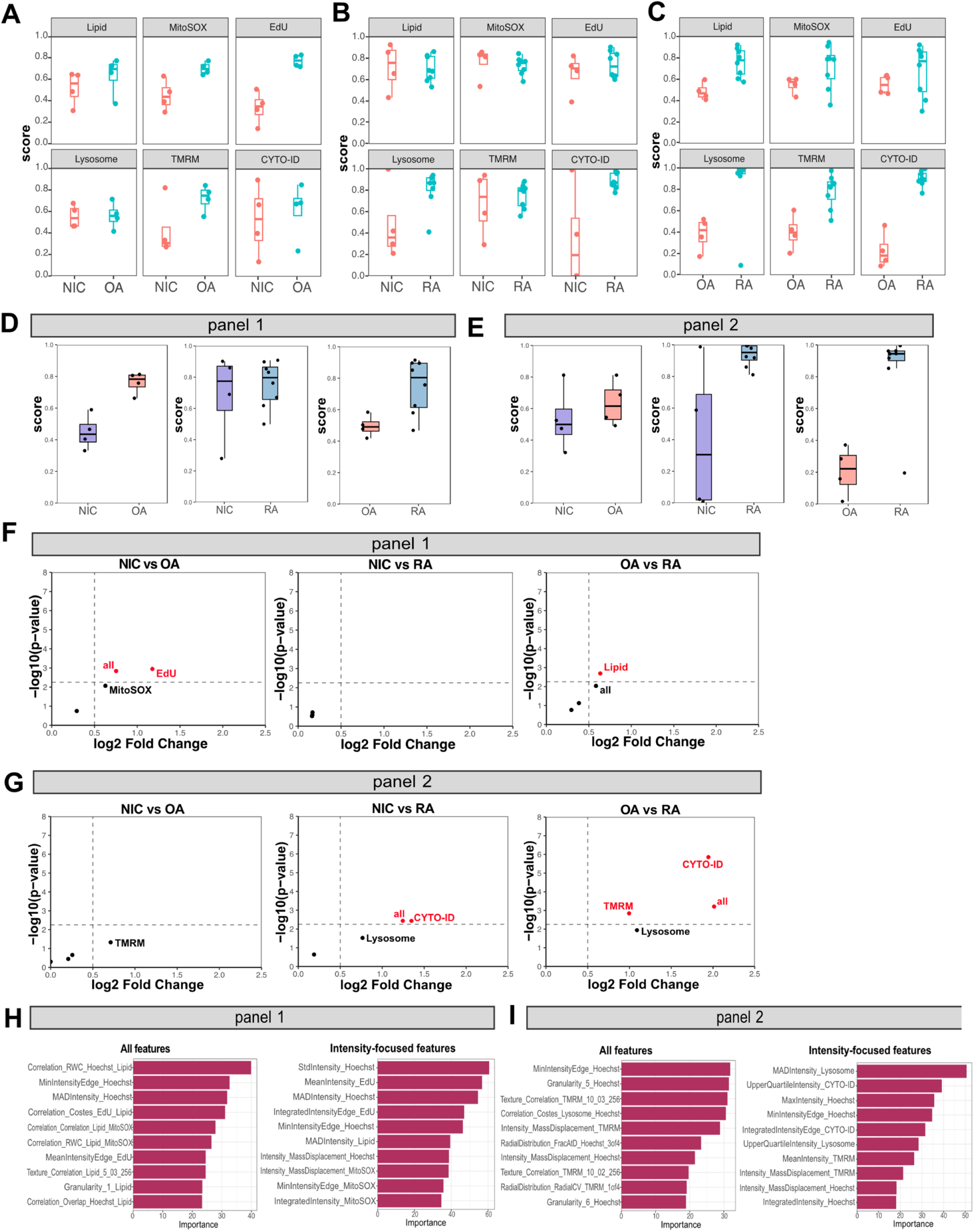
Machine learning classification to distinguish disease groups across multidimensional feature sets. Random forest classification with leave-one-out cross-validation (LOO) was applied to discriminate disease groups using control samples only. (A–C) LOO plots per functional readout comparing (A) NIC vs. OA, (B) NIC vs. RA, and (C) OA vs. RA. (D–E) LOO plots using all markers from panel 1 (D) and panel 2 (E). (F–G) Volcano plots summarizing pairwise log fold changes and LOO p-values from panel 1 (F) and panel 2 (G). (H–I) Feature importance analysis for panel 1 (H) and panel 2 (I), performed in untreated control conditions to identify features that best discriminate between NIC (n = 4), OA (n = 4), and RA (n = 8) groups. To reduce multicollinearity, representative features were selected following correlation clustering and VIF filtering (VIF < 10); importance scores reflect mean decrease in accuracy upon feature permutation. Left bar plots show all representative features; right plots highlight intensity-focused features ranked by importance.

To determine whether integrating all functional readouts into a composite score improves disease discrimination over individual markers, volcano plots were constructed for each pairwise group comparison across both imaging panels (Fig. 2F–G). When applying a combined significance and fold change threshold, only the composite score (“all”) and EdU emerged as statistically robust discriminators in panel 1, with EdU driving the NIC vs OA separation and lipid content distinguishing OA from RA (Fig. 2F). In panel 2, CYTO-ID was the strongest individual discriminator, particularly for NIC vs RA and OA vs RA comparisons, with TMRM and lysosomal content reaching significance only in the OA vs RA contrast (Fig. 2G). Notably, MitoSOX and lysosome— despite contributing to disease separation in multidimensional analysis — did not reach significance as a standalone feature in this volcano framework, highlighting the added sensitivity of integrating multiple readouts. Critically, the composite score (“all”) performed comparably to or exceeded the most informative individual markers across comparisons, supporting the value of a multi-feature approach for capturing the full spectrum of functional differences between disease states.

To address multicollinearity, hierarchical clustering of Pearson correlations was applied to select representative non-redundant discriminative features, confirming consistent results with the findings above. The most discriminatory features spanned multiple biological dimensions: spatial distribution and co-localization metrics of neutral lipid content (Pearson, rank-weighted colocalization [RWC], and Costes coefficients), peripheral EdU intensity, texture and intensity features of TMRM reflecting mitochondrial membrane potential heterogeneity, and intensity-based metrics of CYTO-ID and Lysosome (Fig. 2H–I).

Together, these findings demonstrate that complex multivariate imaging features robustly separate NIC, OA, and RA fibroblasts, capturing disease-specific phenotypes beyond basic intensity measurements.

### Assessment of Synovial Fibroblast Responses to Inflammatory Cytokine Stimulation

To further confirm that our assays could robustly capture biologically meaningful responses, we treated SFs with a range of inflammatory cytokines and immune stimuli: TNF, IFNƔ, IL-1β, the TLR4 ligand LPS, and the TLR3 ligand Poly I:C (PIC). Cytokine treatment induced several stimulus-specific functional changes (mean, SD and p-values are presented in Supplementary Table 1). TNF stimulation increased proliferation across all disease groups, most prominently in OA and RA fibroblasts, consistent with previous data (21) (Fig. 3A, Supplementary Fig. 3A,). Conversely, IFNƔ and PIC suppressed proliferation, with the most pronounced inhibitory effect in RA (Fig. 3A).

**Fig. 3.**
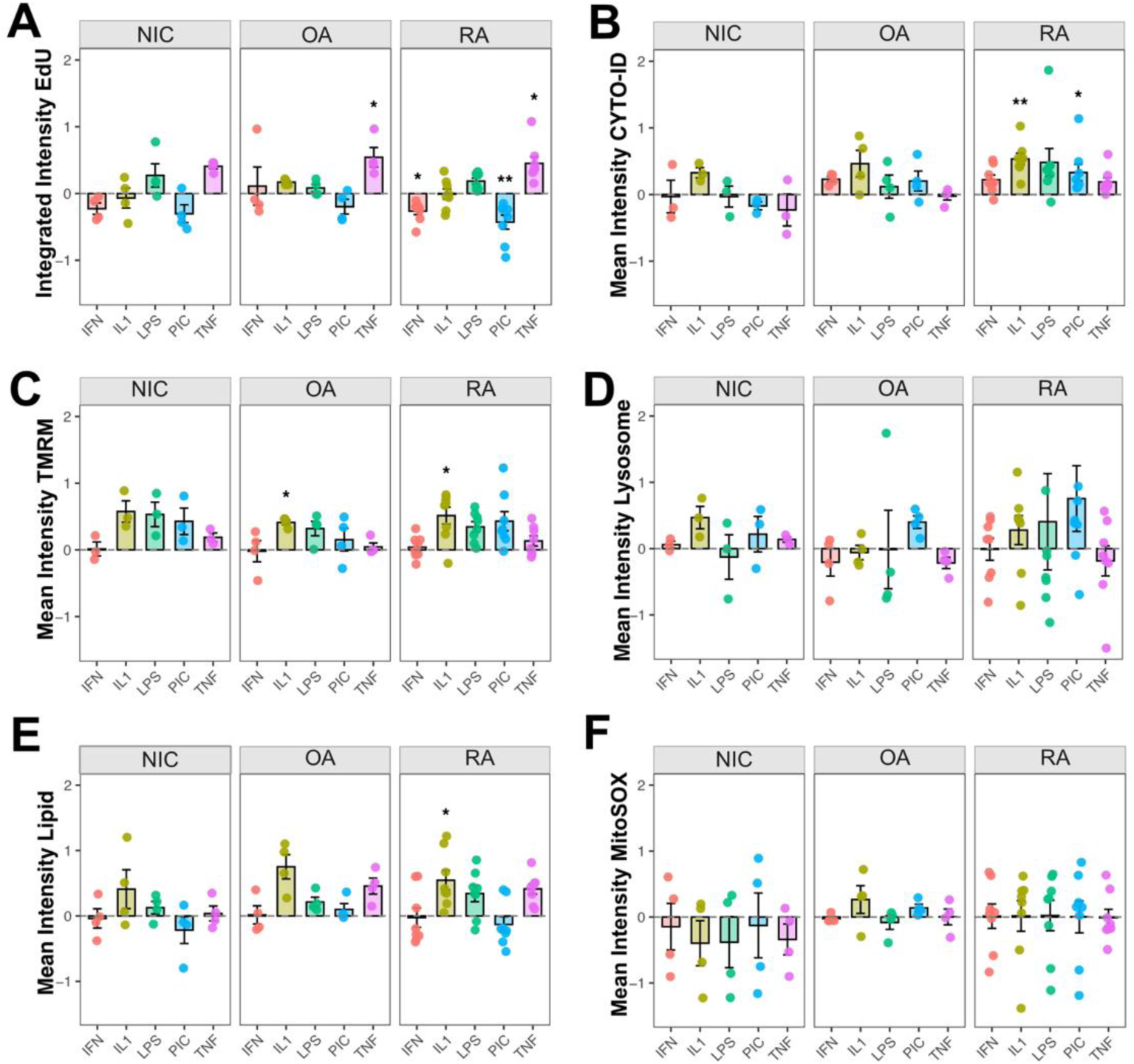
Synovial fibroblast functional responses to inflammatory cytokine stimulation. Bar plots show the fold change in intensity features upon cytokine/stimulant treatment with IFNƔ (IFN), IL-1β (IL1), TNF (TNF), LPS, and Poly I:C (PIC) relative to untreated controls, separately for NIC (n = 4), OA (n = 4), and RA (n = 8) groups. Functional readouts include fold change of (A) integrated intensity of EdU-positive cells (proliferation); (B) CYTO-ID mean intensity (autophagosome); (C) TMRM mean intensity (mitochondrial membrane potential); (D) Lysosome mean intensity; (E) LipidTOX mean intensity (neutral lipid content); (F) MitoSOX mean intensity (mitochondrial superoxide). Each dot represents an individual patient. Pairwise Wilcoxon tests versus control; ** p.adj ≤ 0.01, * p.adj ≤ 0.05.

IL-1β treatment elevated both autophagic activity and mitochondrial membrane potential, with effects most pronounced in RA SFs (Fig. 3B–C). PIC also showed trends toward increased autophagosome content and mitochondrial membrane potential across fibroblast groups, though some did not reach statistical significance (Fig. 3B–C). PIC was the strongest inducer of lysosomal content, with the strongest, but non-significant trend observed in RA SFs (Fig. 3D). This effect of PIC stimulation is potentially linked to the requirement of an acidic environment for its activation (22). In addition to its effects on autophagosomes and mitochondria, IL-1β induced substantial lipid accumulation in OA and RA fibroblasts, with a significant increase in RA (Fig. 3E). Mitochondrial superoxide levels remained unchanged across all pro-inflammatory treatments (Fig. 3F).

Together, these findings demonstrate the capacity of our high-content assays to detect differential, phenotype-driven responses of SFs to inflammatory stimulation. Most notable, was the strong and consistent effect of IL-1β across most of the measured parameters.

### IL-1β Induces Disease-Specific Lipid Redistribution in SFs

To further investigate the functionally relevant imaging features associated with IL-1β-treated SFs, we performed a feature importance analysis within each panel. This analysis identified lipid-related features as the most prominent IL-1β-induced alterations in panel 1 (Fig. 4A). IL-1β stimulation increased peripheral lipid accumulation across all disease groups, reflected by elevated MaxIntensityEdge (Supplementary Fig. 4A). However, granularity texture analysis, (Granularity_3_Lipid, which reflects the proportion of small-to-intermediate sized lipid structures) showed the most pronounced increase in punctate lipid droplet distribution following IL-1β treatment in RA SFs (Supplementary Fig. 4B), suggesting a disease-specific shift toward a more punctate lipid droplet distribution under inflammatory stress. Furthermore, Correlation_Costes_Lipid_EdU decreased following IL-1β treatment (Supplementary Fig. 4C), indicating that lipid droplets particularly dissociate from perinuclear areas in EdU negative cells. This is consistent with their peripheral redistribution observed in MaxIntensityEdge and suggests a divergence between lipid-accumulating and actively proliferating cell states under inflammatory stress. Focusing our analysis on lipid-associated features (Fig. 4B) confirmed spatial lipid redistribution with most feature changes after IL-1β treatment. Again, peripheral redistribution of lipid droplets reflected by increased RadialCV_Lipid 3of4, increased specifically in RA (Supplementary Fig. 4D-E). Notably, RA SFs exhibited visibly distinct lipid patterns under IL-1β compared to NIC SFs (Fig. 4C).

**Fig. 4.**
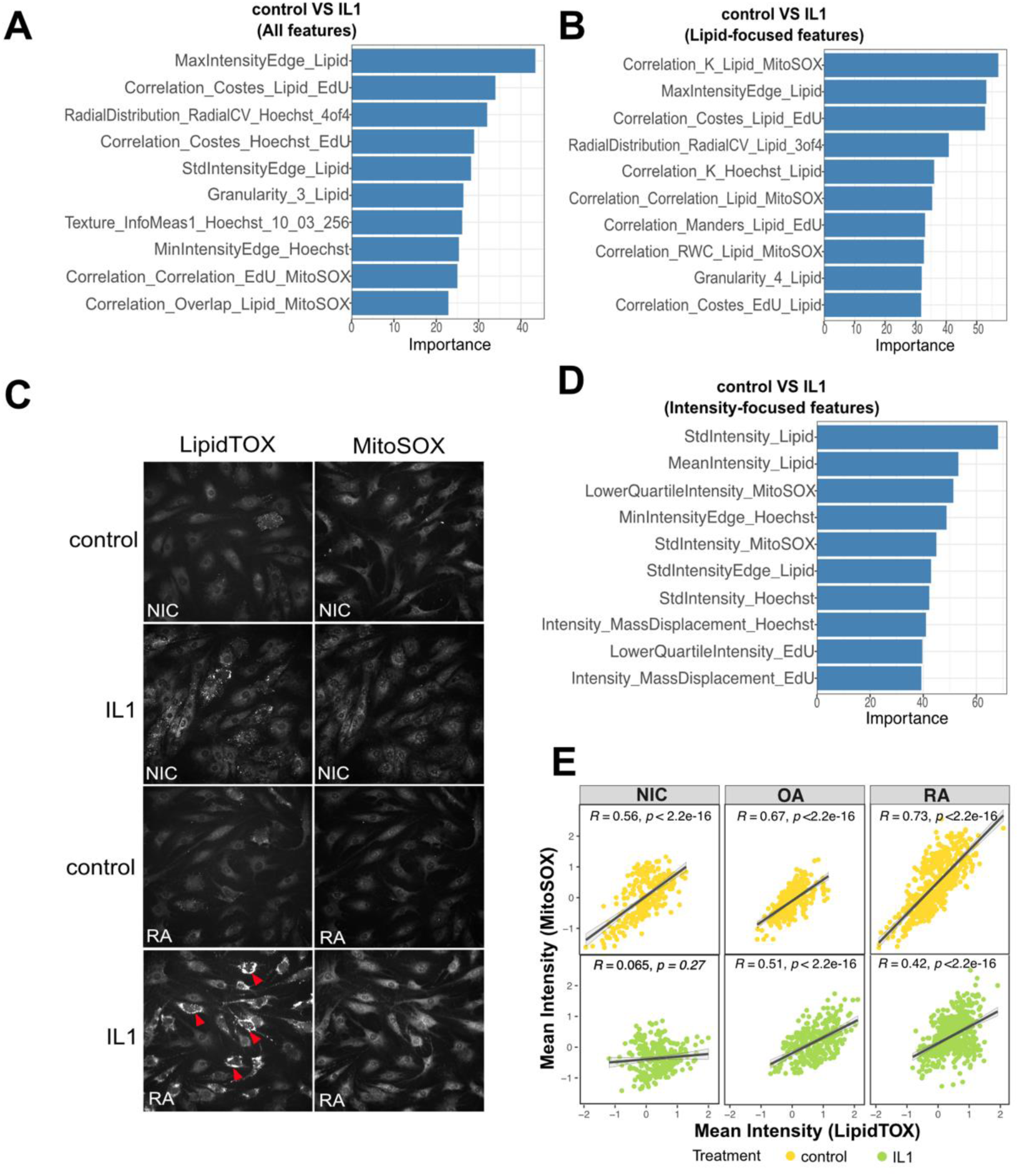
Feature importance analysis reveals IL-1β-induced alterations in lipid metabolism. Random forest feature importance (ranked by MeanDecreaseAccuracy) comparing IL-1β-treated vs. control SFs across all disease groups (NIC; n = 4, OA; n = 4, and RA; n = 8). (A) All features from the metabolism-centered panel. (B) Lipid-focused feature subset. (C) Representative LipidTOX and MitoSOX images comparing control and IL-1β-treated NIC (n = 4) and RA (n = 8) fibroblasts, illustrating disease-specific lipid redistribution patterns particularly in RA. (D) Intensity-focused feature importance following IL-1β treatment. (E) Intercellular Spearman correlation of mean LipidTOX and MitoSOX fluorescence intensities across NIC, OA, and RA fibroblasts, with regression lines and 95% confidence intervals overlaid.

Lipid-focused (Fig. 4B) as well as intensity focused feature analysis (Fig. 4D) highlighted changing correlations between neutral lipid and MitoSOX. Correlation of mean intensity values showed a strong positive correlation between LipidTOX and MitoSOX intensity in unstimulated SFs, which became weaker in OA and RA and was lost in NIC SFs after stimulation with IL-1β (Fig. 4E). This indicates that mitochondrial oxidative stress and lipid accumulation are linked processes in steady state conditions, but become decoupled after stimulation.

Together, these findings suggest that IL-1β reorganizes lipid distribution in fibroblasts and decouples lipid metabolism from both mitochondrial ROS production and proliferation zones. These changes were more pronounced in RA, indicative of disease-specific metabolic stress responses.

### Feature Importance Analysis Reveals IL-1β And PIC Drive Distinct Subcellular Autophagy Signatures in SFs

Further analysis of panel 2 features showed that IL-1β enhanced intracellular spatial co-localization between mitochondria and autophagosomes (RWC of TMRM and CYTO-ID), indicative of mitophagy-like processes (Fig. 5A). Interestingly, this co-localization was much less in NIC SF and was also not significantly increased after stimulation in these cells. TMRM and CYTO-ID co-localization was similarly induced in MG132 treated SFs but distinct in R+C-induced autophagy, where its direction was opposite (Fig. 5B). Notably, the correlation between population-wide intensity signals of TMRM and CYTO-ID was consistent for both IL1 and R+C treatments (Supplementary Fig. 5A–B), suggesting that while overall cellular stress increases similarly, the spatial organization of the stress response depends on the specific trigger.

**Fig. 5.**
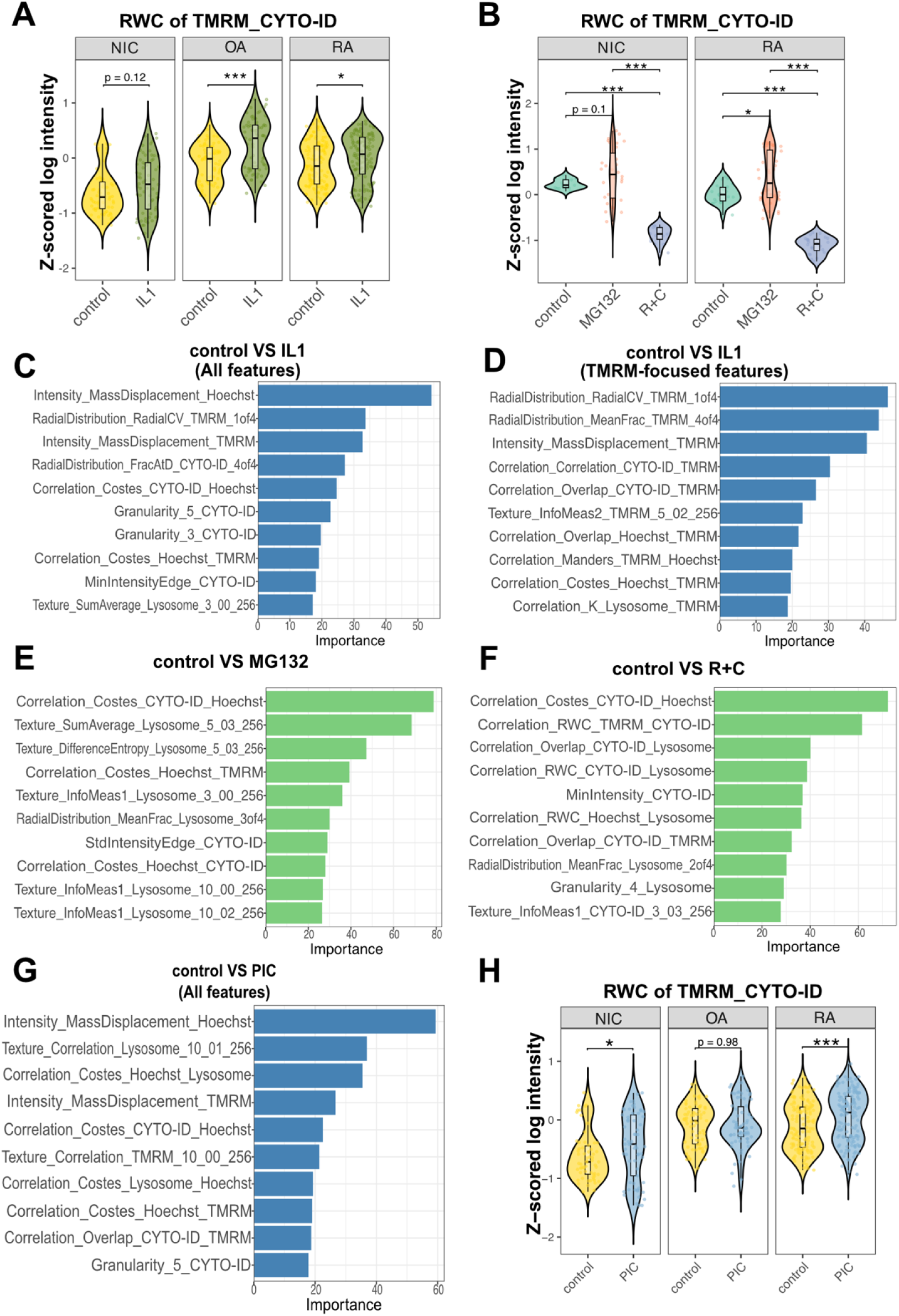
Feature importance analysis linking cytokine treatment to autophagy-related cellular changes. (A–B) violin plots of TMRM–CYTO-ID Rank Weighted Colocalization (RWC) following IL-1β treatment or positive controls (MG132, R+C; n=2). Random forest feature importance (ranked by MeanDecreaseAccuracy) for pairwise comparisons against control across all diseases. (C) All extracted features comparing IL-1β-treated vs. control fibroblasts across all disease groups, highlighting the strongest discriminators of IL-1β stimulation. (D) TMRM-focused subset of the same comparison, illustrating the prominent contribution of mitochondrial distribution and intensity parameters. (E–F) Feature importance for positive controls: R+C and MG132, showing distinct signatures associated with canonical autophagy induction and proteasome inhibition respectively. (G) All features comparing PIC-treated vs. control fibroblasts across all disease groups, highlighting the contribution lysosomal metrics to the PIC response. (H) violin plots of TMRM–CYTO-ID RWC co-localization following PIC treatment.

Feature importance analysis compared to controls within panel 2, revealed that IL-1β treatment mostly affected nuclear, TMRM and CYTO-ID features. It shifted mitochondrial activity in both subcellular localization and distribution, alongside altered nuclear staining patterns (Fig. 5C). Elevated Mass Displacement metrics indicated peripheral polarization of both nuclear and mitochondrial signals, suggesting spatial reorganization under inflammatory stress (Supplementary Fig. 5C–D). Radial distribution parameters — particularly RadialDistribution_RadialCV_TMRM_1of4 — further reflected heterogeneous perinuclear mitochondrial clustering in response to IL-1β (Fig. 5C, Supplementary Fig. 5E). The prominence of TMRM-related features confirmed the findings of the global panel analysis, with mitochondrial membrane potential changes correlating with the autophagy marker CYTO-ID (Fig. 5D). In contrast, MG132 and R+C treatment drove markedly different feature profiles including several changes in lysosomes (Fig. 5E–F), supportive of differential autophagic engagement. While IL-1β appeared to preferentially involve mitophagy-like processes, R+C drove broader autophagosome accumulation.

IL-1β and PIC stimulation both induced autophagic and mitochondrial membrane potential responses (Fig. 3B–C); however, deeper feature-level analysis revealed both shared and divergent patterns. Similar to IL-1β and MG132, TMRM and Hoechst MassDisplacement was also elevated after PIC (Fig. 5C, G; Supplementary Fig. 5C-D, F-H), consistent with stress-induced mitochondrial redistribution and cytoskeletal reorganization. Mitochondria-autophagosome co-localization (RWC of TMRM and CYTO-ID) was also similarly increased under both IL-1β and PIC. However, PIC induced this response in NIC and RA SFs, but not in OA cells (Fig. 5H). Despite these shared features, the two conditions diverged in their lysosomal engagement: PIC uniquely drove prominent lysosome-related texture changes, particularly in RA SFs (Fig. 5G; Supplementary Fig. 5I), suggesting preferential activation of lysosomal-autophagic flux, in contrast to the mitochondria-centered autophagic response characteristic of IL-1β stimulation.

### Identification of candidate genes underlying IL-1β-Triggered Autophagy

To link our functional imaging observations with molecular mechanisms, we analyzed a publicly available RNA sequencing dataset from Tsuchiya *et al.* (23), in which RA and OA SFs were stimulated with inflammatory cytokines. We mapped autophagy-related genes using the curated Human Autophagy Database (24).

In gene set enrichment analysis (GSEA), IL-1β stimulation significantly upregulated the annotated autophagy gene set in SFs (Fig. 6A). From the 25 leading-edge genes, we identified 12 with established roles in autophagosome formation and dynamics (marked bold in Fig. 6B). Tracing the autophagic cascade, *HIF1A, EPAS1*, and *BNIP3* represent a coordinated hypoxia-sensing axis, wherein HIF-1α and HIF-2α (encoded by *EPAS1*) transcriptionally induce BNIP3-mediated mitophagy under oxidative and hypoxic stress (25,26); RCAN1 further reinforces this mitophagy response as an additional stress-responsive regulator (27,27), together consistent with the mitochondrial–autophagosome co-localization observed in Fig. 5A. PIM2 and TRIM16 coordinate upstream machinery assembly (28,29), while WIPI1 drives phagophore nucleation at PI3P-enriched membranes (30). ATG7 and TP53INP2 cooperate in LC3 lipidation and membrane conjugation (1), and ATG4D–GABARAPL1 function as a linked pair mediating cargo tethering and autophagosome–lysosome fusion (33,34). DRAM1 completes the process by facilitating autophagosome–lysosome fusion (35).

**Fig. 6.**
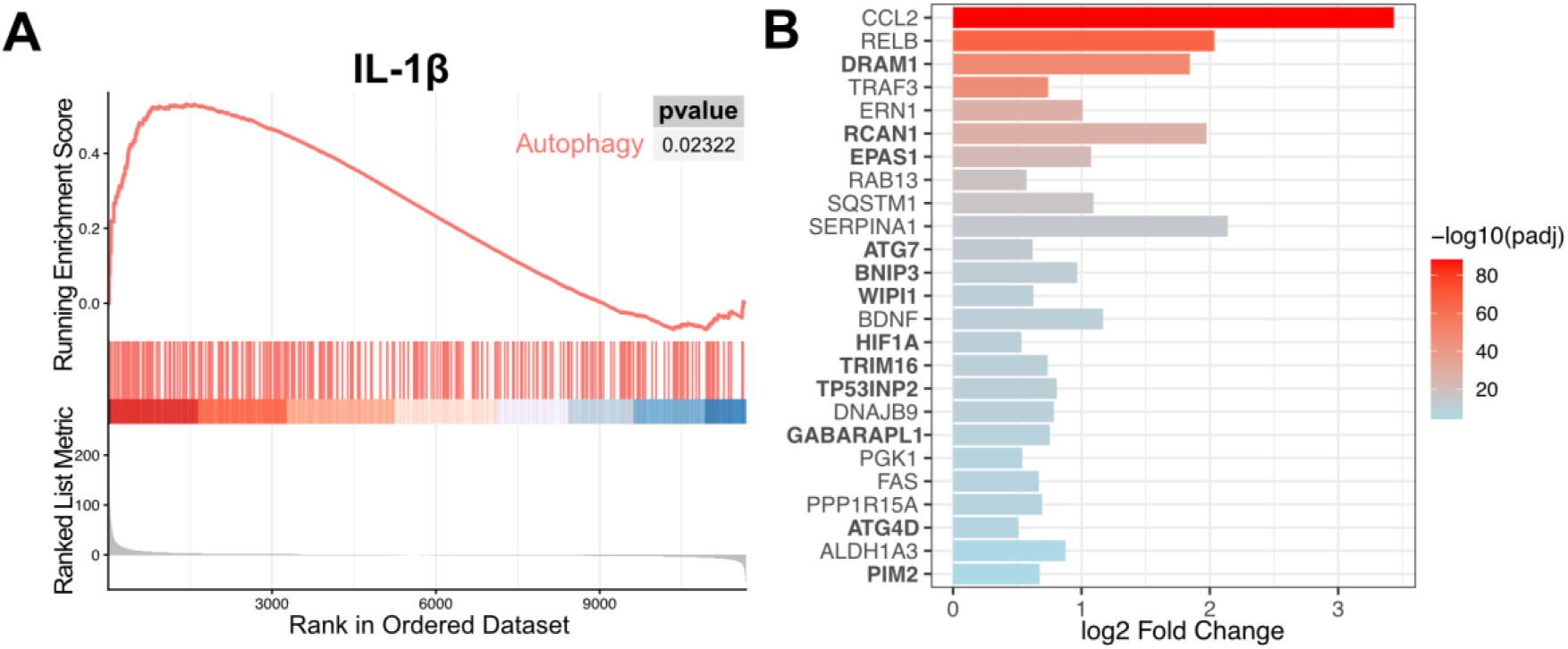
GSEA of autophagy-related genes in cytokine-stimulated synovial fibroblasts. Data were derived from differential expression analysis of cytokine-stimulated synovial fibroblasts (23). (A) GSEA enrichment plots for the autophagy gene set (curated from HADb (24)) in RA and OA fibroblasts stimulated with IL-1β, showing normalized enrichment scores (NES), p-value suggesting positive enrichment. (B) Bar plot of the 25 leading-edge genes upregulated by IL-1β stimulation, ordered by -log10(padj) Color intensity reflects −log10 adjusted p-value. Genes with established roles in autophagosome formation and dynamics are highlighted.

Collectively, this transcriptomic analysis provides evidence that IL-1β potently activates a coordinated autophagy program in SFs, with particular enrichment of the HIF-1α–BNIP3 mitophagy axis.

## Discussion

In this study, we developed two high-content functional panels targeting autophagy dynamics, mitochondrial health, lipid metabolism, and proliferation to comprehensively capture disease relevant functional changes in SFs. We first extracted commonly used intensity-based features before applying machine learning to leverage the full dimensionality of imaging data for classification and feature discovery. We found that texture and spatial metrics proved particularly informative to reveal stimulation and disease-specific differences, illustrating how in-depth feature analysis in high content imaging can reveal distributional differences invisible to mean intensity alone. Feature importance analysis uncovered spatial relationships and cellular heterogeneity beyond conventional intensity metrics, providing a rich feature set for understanding fibroblast phenotypes.

We show that mitochondrial health, lipid metabolism and autophagy related features were most useful in differentiation of unstimulated RA SFs from OA SFs. OA SFs exhibited higher basal autophagy activity than RA, contrasting with previous reports of increased autophagy in RA compared to OA SFs. However, our data might reflect an imprinted disturbance in steady state autophagosome formation in RA SFs that might or might not influence the autophagic flux in RA SFs when autophagy is activated (36). CYTO-ID was selected for its high-throughput compatibility and multiplex suitability, though it cannot independently distinguish increased autophagosome formation from impaired lysosomal degradation; complementary markers and pharmacological flux assays would be required for precise flux assessment.

We confirmed biologically relevant assay responses by cytokine stimulations. Even though samples numbers in our study were too small to robustly capture differences in response between NIC, OA and RA SFs, the heterogenous response patterns that we observed point to altered inflammatory response pathways in RA as well as OA SFs. Among all stimuli, IL-1β elicited the most pronounced effects on autophagy, mitochondrial membrane potential, and lipid metabolism. Since these functional changes were also discriminative between RA and OA SFs in basal conditions, it might be speculated that elevated IL-1 β in RA synovial tissue and fluid might lead to persistent epigenetic modulations in these functions in RA SFs (37,38).

Feature importance analysis identified IL-1β-induced neutral lipid accumulation particularly in RA SFs, with spatial and textural changes consistent with raw image staining patterns. Differences in lipid content have been previously suggested in RA SFs (10). However, up to now, these findings remain exploratory, and dedicated validation, e.g. with flux blockers, genetic knockdowns, and orthogonal readouts, will be essential to confirm mechanistic interpretations. Disease-specific alterations in lipid production, transport, or clearance are a promising new area for further investigation in RA SFs.

Our data indicate that IL-1β stimulation drives a coordinated, selective autophagy transcriptional program in SFs. The IL-1β-driven autophagy gene signature that we identified suggests that IL-1β promotes mitochondria-targeted autophagy through a HIF-1α–BNIP3 pseudo-hypoxic axis (25,26), which is consistent with the mitophagy-like spatial redistribution observed in our imaging data. Particularly notable is the upregulation of *RCAN1*, a gene whose risk variants have been mapped to SF-specific enhancers in a functional genomics atlas of RA heritability (39), implicating it as an SF-intrinsic RA risk gene. Beyond its genetic relevance, RCAN1 is a stress-inducible inhibitor of calcineurin that regulates the calcineurin–NFATc1 pathway governing osteoclastogenesis and bone homeostasis and has been linked to mitophagy activation under oxidative and hypoxic stress (27,40,41), directly converging with the mitochondrial autophagy phenotype observed here. Together, our findings provide convergent evidence from functional imaging and transcriptomic analysis that IL-1β drives an adaptive mitophagy program in RA SFs, with RCAN1 emerging as a mechanistically and genetically relevant mediator. Direct causal relationships, however, remain to be established through targeted functional experiments.

Several limitations warrant consideration. Channel constraints required reserving the 488 nm channel exclusively for CYTO-ID, limiting multiplexing flexibility. Cytoplasm segmentation relied on broadly distributed marker staining rather than a dedicated membrane dye, a trade-off between channel efficiency and segmentation precision addressable in future iterations. Disease comparisons should be interpreted cautiously given sample size limitations; cohort expansion and orthogonal assay integration will be necessary to refine findings. When scaling across plates or batches, careful experimental design — including randomized sample allocation, bridging controls, and donor-matched alignment — is critical to preserve biological signal. Finally, direct causal relationships between IL-1β signaling, the identified driver genes, and the autophagy phenotype observed in SFs remain to be established through targeted functional experiments.

For the first time, our study combines the measurement of six different functional readouts to characterize the basal and activated phenotypes of NIC, OA and RA SFs. Our findings underscore the value of integrating high-content image analysis with functional assay platforms. Our approach is readily applicable to drug screening, genetic perturbation studies, and broader experimental contexts beyond cytokine treatment.

## Funding

This work was funded by the Swiss National Research Foundation (Grant Number 310030_215118.

## Acknowledgements

We thank Peter Künzler for excellent technical support in culturing cells. We thank Kristina Bürki, Raphael Micheroli, Katharina Zachariassen, Thomas Rauer, Esin Rothenfluh, and Miriam Marks for organizing synovial tissues from patients and controls.

## Notes

### Competing Interest Statement

The authors have declared no competing interest.

