## Supplementary Figures 1-5, Supplementary Tables for "Developing High Content Imaging Functional Panels to Characterize Synovial Fibroblasts in Rheumatoid Arthritis"

#### Supplementary Figure 1

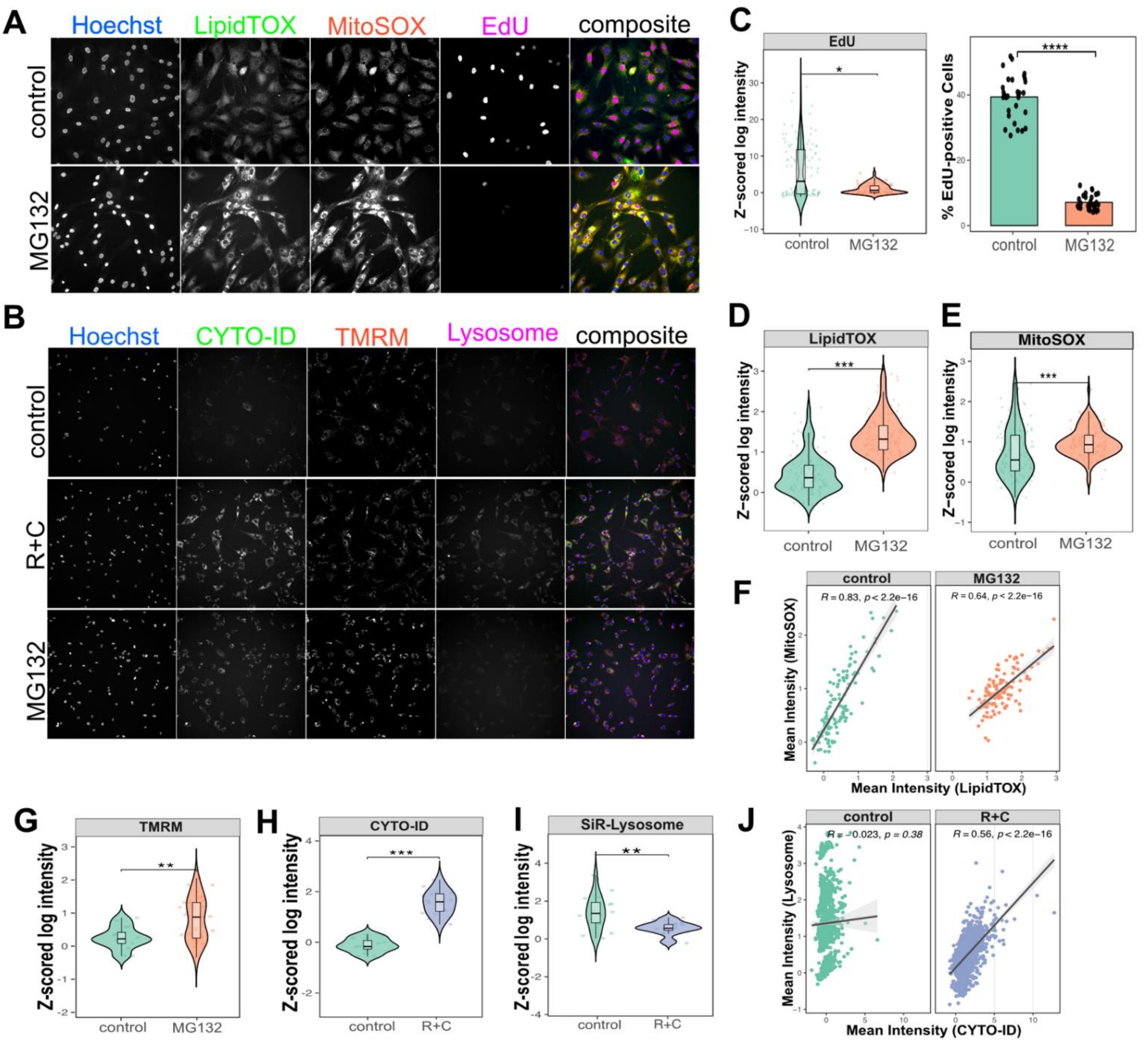


**Supplementary Fig. 1 Functional panel development and validation.**
Assays were performed on non-inflammatory control synovial fibroblasts (NIC SFs; n=2). Representative images from (A) panel 1 showing Hoechst, LipidTOX, MitoSOX, EdU, and composite overlay, with MG132 as positive control, and (B) panel 2 showing Hoechst, CYTO-ID, TMRM, SiR-Lysosome, and composite overlay, with MG132 or rapamycine plus chloroquine (R+C) as positive controls. (C) Violin plots of EdU mean intensity and percentage EdU-positive cells per field following MG132 treatment. Violin plots of mean intensities for (D) LipidTOX, (E) MitoSOX, (G) TMRM, (H) CYTO-ID, and (I) SiR-Lysosome after positive control treatment. Scatter plots showing Spearman correlations between (F) LipidTOX and MitoSOX and (J) CYTO-ID and SiR-Lysosome intensities. Statistical comparisons used Wilcoxon tests with Bonferroni-adjusted p-values (**** p.adj ≤ 0.0001, *** p.adj ≤ 0.001, ** p.adj ≤ 0.01) and Spearman correlation for (F), (J).

#### Supplementary Figure 2

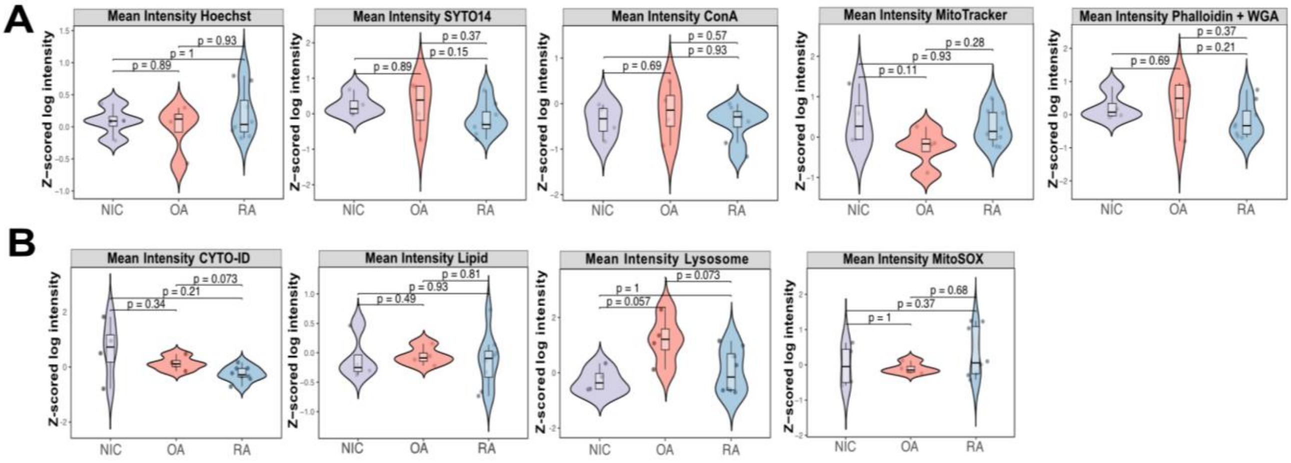


**Supplementary Fig. 2 Disease-associated differences in functional profiles of synovial fibroblasts.** Violin plots of per-patient mean fluorescence intensity distributions at baseline. (A) Morphological markers from an adapted Cell Painting assay from Carpenter et al. (17): Hoechst (nucleus), SYTO (RNA/nucleoli), ConA (ER/Golgi), MitoTracker (mitochondria), and Phalloidin + WGA (actin/Golgi/plasma membrane). (B) Functional markers: CYTO-ID (autophagosomes), LipidTOX (neutral lipids), Lysosome, and MitoSOX (mitochondrial superoxide). No significant differences were detected between NIC, OA, and RA fibroblasts by pairwise Wilcoxon rank-sum tests.

**Supplementary Figure 3**

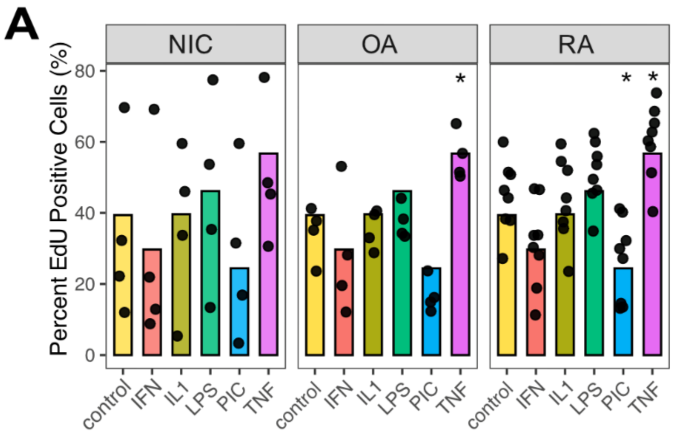


**Supplementary Fig. 3 Increased proliferation marker after TNFα stimulation.**
(A) Bar plots showed percent EdU positive cell after cytokine stimulation; IFNɣ (IFN), IL-1β (IL1), TNFα (TNF), LPS, and Poly I:C (PIC). Each dot represent patient.

**Supplementary Figure 4**

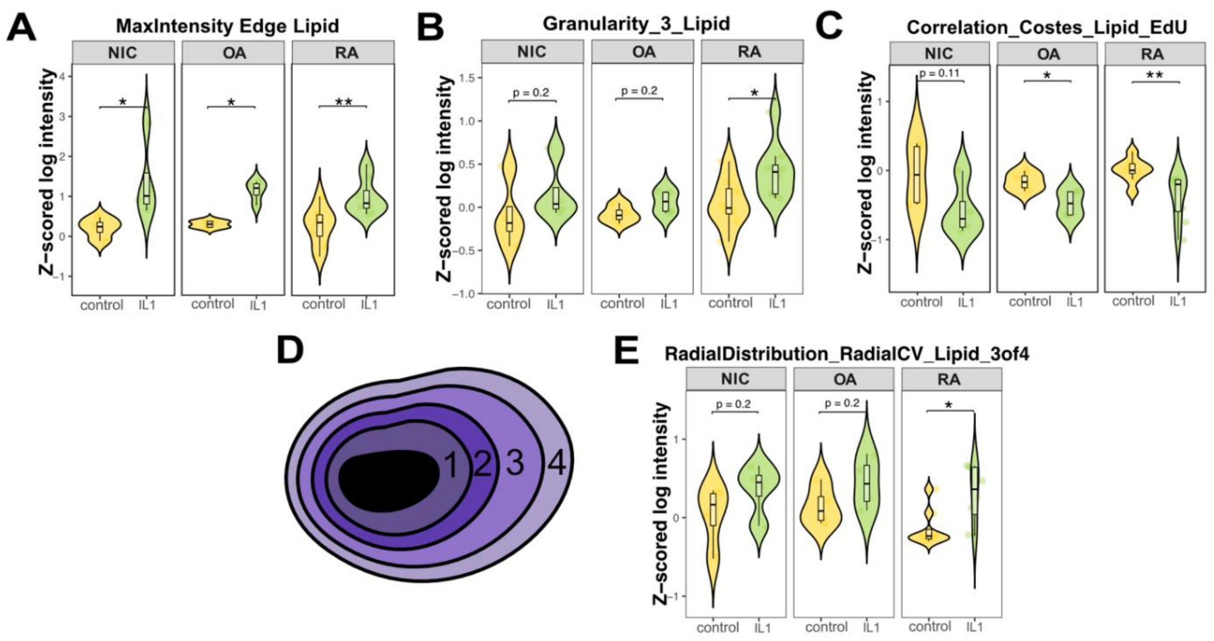


**Supplementary Fig. 4 IL-1β -induced alterations in lipid distribution features across disease groups.**

Violin plots of per-patient distributions comparing NIC, OA, and RA fibroblasts following IL-1β treatment: (A) MaxIntensityEdge of LipidTOX; (B) Granularity_3 of LipidTOX; (C) Costes co-localization coefficient of LipidTOX and EdU; (D) schematic illustrating RadialDistribution measurement zones; (E) RadialCV of LipidTOX 3of4. Statistical comparisons used pairwise Wilcoxon tests with Bonferroni-adjusted p-values (** p.adj ≤ 0.01, * p.adj ≤ 0.05).

**Supplementary Figure 5**

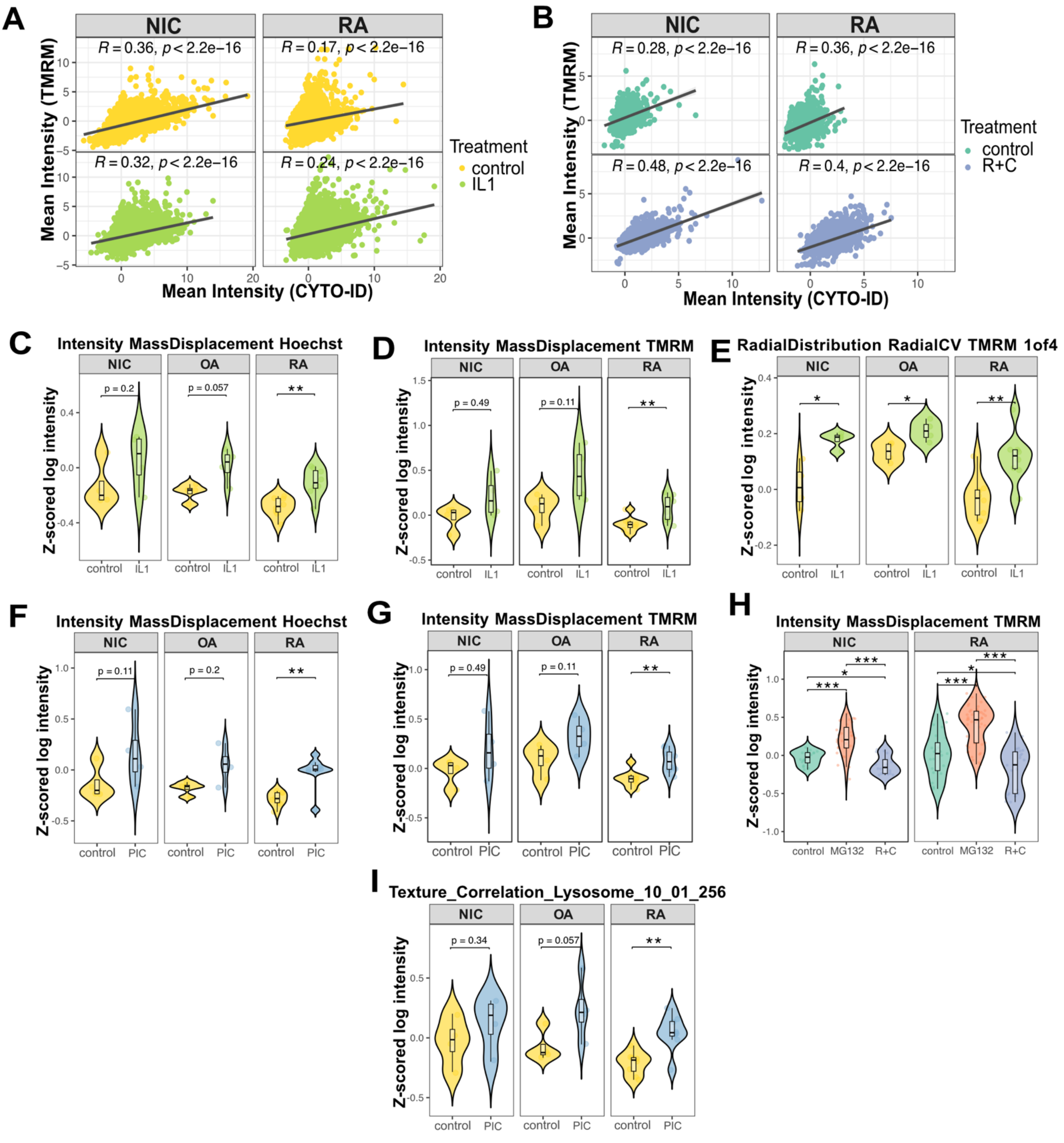


**Supplementary Fig. 5 Intracellular marker redistribution following IL-1β and PIC treatment.**
(A) Scatter plots showing Spearman correlations between mean TMRM and CYTO-ID fluorescence intensities following (A) IL-1β and (B) R+C treatment, with linear regression lines, 95% confidence intervals, and Spearman correlation coefficients and p-values overlaid.
Violin plots of per-patient fluorescence intensity distributions comparing NIC, OA, and RA fibroblasts after IL-1β or PIC treatment. (C, F) MassDisplacement of Hoechst; (D, G) MassDisplacement of TMRM; (E) RadialCV of TMRM 1of4; (H) per-cell MassDisplacement of TMRM after positive controls (MG132, R+C) treatment; (I) Texture Correlation of Lysosome (10_01_256). Statistical comparisons used pairwise Wilcoxon tests with Bonferroni-adjusted p-values (*** p.adj ≤ 0.001, ** p.adj ≤ 0.01, * p.adj ≤ 0.05).

### Supplementary tables

**Supplementary Tables 1–2 Per-patient mean fluorescence intensities for panel 1 and panel 2 features.** Tables 1 and 2 report per-patient mean fluorescence intensities for panel 1 and panel 2, respectively. Statistical comparisons versus control used pairwise Wilcoxon tests on patient-level means (*** p ≤ 0.001, ** p ≤ 0.01, * p ≤ 0.05, ns = not significant).

#### Table 1; Mean intensity of panel 1 after cytokine treatment

| | | Treatment (mean, SD, pvalue, significance) | | | | |
|---|---|---|---|---|---|---|
| Disease | Features | IFN | IL1 | LPS | PIC | TNF |
| NIC | Mean Intensity EdU | mean = 2.208, SD = 2.776 p = 0.486, ns | mean = 2.426, SD = 1.862 p = 1, ns | mean = 4.147, SD = 3.329 p = 0.886, ns | mean = 2.269, SD = 2.18 p = 0.886, ns | mean = 5, SD = 3.749 p = 0.343, ns |
| | Mean Intensity Lipid | mean = -0.102, SD = 0.192 p = 0.886, ns | mean = 0.345, SD = 0.416 p = 0.2, ns | mean = 0.063, SD = 0.233 p = 0.486, ns | mean = -0.281, SD = 0.099 p = 0.343, ns | mean = -0.027, SD = 0.353 p = 0.886, ns |
| | Mean Intensity MitoSOX | mean = -0.124, SD = 0.288 p = 0.886, ns | mean = -0.375, SD = 0.342 p = 0.686, ns | mean = -0.36, SD = 0.318 p = 0.686, ns | mean = -0.105, SD = 0.505 p = 0.886, ns | mean = -0.317, SD = 0.311 p = 0.686, ns |
| OA | Mean Intensity EdU | mean = 1.564, SD = 2.067 p = 0.486, ns | mean = 1.15, SD = 0.2 p = 1, ns | mean = 1.193, SD = 0.127 p = 0.686, ns | mean = 0.432, SD = 0.193 p = 0.029, * | mean = 2.68, SD = 1.196 p = 0.029, * |
| | Mean Intensity Lipid | mean = -0.041, SD = 0.404 p = 0.686, ns | mean = 0.695, SD = 0.437 p = 0.057, ns | mean = 0.156, SD = 0.309 p = 0.486, ns | mean = 0.043, SD = 0.335 p = 0.886, ns | mean = 0.4, SD = 0.4 p = 0.114, ns |
| | Mean Intensity MitoSOX | mean = -0.125, SD = 0.186 p = 0.886, ns | mean = 0.163, SD = 0.321 p = 0.343, ns | mean = -0.184, SD = 0.057 p = 0.686, ns | mean = 0.037, SD = 0.233 p = 0.343, ns | mean = -0.1, SD = 0.145 p = 1, ns |
| RA | Mean Intensity EdU | mean = 1.573, SD = 0.757 p = 0.028, * | mean = 2.452, SD = 1.784 p = 0.382, ns | mean = 3.314, SD = 1.278 p = 0.442, ns | mean = 1.182, SD = 0.445 p = 0.01, ** | mean = 4.367, SD = 2.909 p = 0.13, ns |
| | Mean Intensity Lipid | mean = -0.133, SD = 0.292 p = 1, ns | mean = 0.438, SD = 0.322 p = 0.021, * | mean = 0.235, SD = 0.383 p = 0.105, ns | mean = -0.238, SD = 0.264 p = 0.574, ns | mean = 0.306, SD = 0.395 p = 0.083, ns |
| | Mean Intensity MitoSOX | mean =0.366, SD = 0.423 p = 0.721, ns | mean = 0.37, SD = 0.396 p = 0.574, ns | mean = 0.377, SD = 0.276 p = 0.574, ns | mean = 0.355, SD = 0.355 p = 0.574, ns | mean = 0.342, SD = 0.549 p = 1, ns |

#### Table 2; Mean intensity of panel 2 after cytokine treatment

| | | Treatment (mean, SD, pvalue, significance) | | | | |
|---|---|---|---|---|---|---|
| Disease | Features | IFN | IL1 | LPS | PIC | TNF |
| NIC | Mean Intensity CYTO-ID | mean = 1.08, SD = 0.375 p = 1, ns | mean = 1.437, SD = 0.549 p = 0.4, ns | mean = 1.079, SD = 0.481 p = 1, ns | mean = 0.94, SD = 0.674 p = 0.7, ns | mean = 0.879, SD = 0.585 p = 0.7, ns |
| | Mean Intensity Lysosome | mean = -0.234, SD = 0.548 p = 0.7, ns | mean = 0.174, SD = 0.335 p = 0.4, ns | mean = -0.42, SD = 0.217 p = 1, ns | mean = -0.076, SD = 0.16 p = 0.7, ns | mean = -0.154, SD = 0.474 p = 0.4, ns |
| | Mean Intensity TMRM | mean = -0.361, SD = 0.478 p = 1, ns | mean = 0.201, SD = 0.308 p = 0.2, ns | mean = 0.157, SD = 0.506 p = 0.2, ns | mean = 0.053, SD = 0.479 p = 0.4, ns | mean = -0.187, SD = 0.272 p = 0.7, ns |
| OA | Mean Intensity CYTO-ID | mean = 0.371, SD = 0.207 p = 0.343, ns | mean = 0.603, SD = 0.227 p = 0.114, ns | mean = 0.258, SD = 0.266 p = 0.486, ns | mean = 0.343, SD = 0.118 p = 0.343, ns | mean = 0.115, SD = 0.171 p = 0.886, ns |
| | Mean Intensity Lysosome | mean = 1.021, SD = 0.572 p = 1, ns | mean = 1.167, SD = 0.911 p = 0.886, ns | mean = 1.212, SD = 1.968 p = 0.686, ns | mean = 1.626, SD = 0.897 p = 0.486, ns | mean = 1.009, SD = 1.022 p = 0.686, ns |
| | Mean Intensity TMRM | mean = 0.239, SD = 0.314 p = 1, ns | mean = 0.671, SD = 0.166 p = 0.029, * | mean = 0.578, SD = 0.236 p = 0.114, ns | mean = 0.412, SD = 0.324 p = 0.343, ns | mean = 0.304, SD = 0.19 p = 0.886, ns |
| RA | Mean Intensity CYTO-ID | mean = 0.003, SD = 0.379 p = 0.328, ns | mean = 0.314, SD = 0.322 p = 0.005, ** | mean = 0.263, SD = 0.452 p = 0.028, * | mean = 0.111, SD = 0.225 p = 0.028, * | mean = -0.032, SD = 0.339 p = 0.234, ns |
| | Mean Intensity Lysosome | mean = 0.038, SD = 0.744 p = 0.878, ns | mean = 0.328, SD = 0.682 p = 0.505, ns | mean = 0.453, SD = 2.466 p = 0.798, ns | mean = 0.804, SD = 1.871 p = 0.505, ns | mean = -0.137, SD = 0.682 p = 0.442, ns |
| | Mean Intensity TMRM | mean = -0.083, SD = 0.369 p = 0.959, ns | mean = 0.39, SD = 0.426 p = 0.038, * | mean = 0.222, SD = 0.397 p = 0.105, ns | mean = 0.307, SD = 0.499 p = 0.083, ns | mean = 0.011, SD = 0.305 p = 0.645, ns |

NOTE: Statistical test (pairwise_wilcox_test) were performed compared to control treatment.
