## Supplementary Methods for "Developing High Content Imaging Functional Panels to Characterize Synovial Fibroblasts in Rheumatoid Arthritis"

***Functional staining protocol***

**Table1 Specification of multiplex staining dyes**

| **Fluorescent dyes (wavelength Ex, Em)** | **Marker** | **Working concentration** | **filters** | **fixation compatibility** | **live staining capability** | **Manufacturer (Cat. No.)** |
| --- | --- | --- | --- | --- | --- | --- |
| **Panel 1** | | | | | | |
| HCS LipidTOX™ Green (495,505 nm) | Neutral lipid | 1:200 | 485-20 BGRFRN_BGRFRN | yes | no | Invitrogen™ (H34475) |
| MitoSOX™ Red (396,610 nm) | Mitochondrial reactive oxygen species (superoxide specifically) | 1:1000 (5 µM) | 549-15 BGRFRN_RS | yes | Label in live cells | Invitrogen (M36007) |
| Click-iT® EdU (5-ethynyl-2'-deoxyuridine) (650, 670 nm) | Proliferation | 10 µM | 650-13 BGRFRN_BGRFRN | yes | Label in live cells | Invitrogen™ (C10340) |
| **Panel 2** | | | | | | |
| CYTO-ID® (480,530 nm) | Autophagic compartments including autophagosomes and autolysosomes. | 1:500 | 485-20 BGRFRN_BGRFRN | yes | yes | Enzo (ENZ-51031) |
| Image-iT™ TMRM Reagent (Tetramethylrhodamine methyl ester) (548,574 nm) | Mitochondrial membrane potential | 100 nM | 549-15 BGRFRN_BGRFRN | no | yes | Invitrogen™ (I34361) |
| SiR-Lysosome (652,674 nm) | Lysosome | 1 µM (with 10 µM Verapamil; efflux pump inhibitor) | 650-13 BGRFRN_BGRFRN | no | yes | Spirochrome (SC012) |
| **Both panel** | | | | | | |
| Hoechst 33342 (340, 480 nm) | Nuclear DNA | 1 µg/mL | 386-23 BGRFRN_BGRFRN | yes | yes | Invitrogen™ (H3570) |

**NOTE: filter from CX7 high content imager**

*Staining steps*

*Panel 1:*

1. Optionally, pretreat cells with positive control compounds or inflammatory cytokines in complete culture medium for 24 hours.
2. Sixteen to eighteen hours before staining, pulse cells with 2× EdU solution prepared in complete medium (added directly to the existing treatment medium) to achieve a final concentration of 10 µM.
3. After 24 hours, remove the medium containing drug, cytokine, and EdU, and add MitoSOX solution in complete medium for live cell labelling.
4. Incubate for 30 minutes at 37°C.
5. Fix cells by adding 16% formaldehyde to reach a final concentration of 3.5–4% and incubate for 20 minutes at room temperature.
6. Remove the fixative and wash once with PBS.
7. Add LipidTOX solution and incubate for 30 minutes at room temperature.
8. Remove LipidTOX and wash once with PBS, then permeabilize cells with 0.1% saponin solution for 15 minutes.
9. Prepare and apply Click-iT reaction solution according to the manufacturer’s instructions and incubate for 30 minutes at room temperature.
10. Wash once with PBS.
11. Stain with Hoechst 33342 solution in PBS for 15 minutes.
12. Wash once with PBS and maintain cells in fresh PBS.
13. Acquire images.

*Panel 2:*

1. Optionally, pretreat cells with positive control compounds or inflammatory cytokines in complete culture medium for 24 hours.
2. Prepare the staining solution by mixing TMRM, CYTO-ID, Lysosome dye, verapamil, and Hoechst 33342 in complete culture medium.
3. Remove the existing medium and add 50 µL of the staining solution per well.
4. Incubate the cells for 30 minutes at 37°C.
5. Wash once with fresh complete medium and replace with 100 µL of complete medium.
6. Acquire images immediately.

Note

Both staining panels were optimized to minimize washing steps, prevent cell detachment, and facilitate high-throughput or large-scale staining experiments.
